# UNBIAS: An attempt to correct abundance bias in 16S sequencing, with limited success

**DOI:** 10.1101/124149

**Authors:** Robert C. Edgar

## Abstract

Next-generation amplicon sequencing of 16S ribosomal RNA is widely used to survey microbial communities. Alpha and beta diversities of these communities are often quantified on the basis of OTU frequencies in the reads. Read abundances are biased by factors including 16S copy number and PCR primer mismatches which can cause the read abundance distribution to diverge substantially from the species abundance distribution. Using mock community tests with species abundances determined independently by shotgun sequencing, I find that 16S amplicon read frequencies have no meaningful correlation with species frequencies (Pearson coefficient *r* close to zero). In addition, I show that that the Jaccard distance between the abundance distributions for reads of replicate samples, which ideally would be zero, is typically ~0.15 with values up to 0.71 for replicates sequenced in different runs. Using simulated communities, I estimate that the average rank of a dominant species in the reads is 3. I describe UNBIAS, a method that attempts to correct for abundance bias due to gene copy number and primer mismatches. I show that UNBIAS can achieve informative, but still poor, correlations (*r* ~0.6) between estimated and true abundances in the idealized case of mock samples where species are well known. However, *r* falls to ~0.4 when the closest reference species have 97% identity and to ~0.2 at 95% identity. This degradation is mostly explained by the increased difficulty in predicting 16S copy number when OTUs have lower similarity with the reference database, as will typically be the case in practice. 16S abundance bias therefore remains an unsolved problem, calling into question the naive use of alpha and beta diversity metrics based on frequency distributions.

## Introduction

Next-generation sequencing has revolutionized the study of microbial communities in environments ranging from the human body (Cho & Blaser 2012; Pflughoeft & Versalovic 2012) to oceans (Moran 2015) and soils (Hartmann et al. 2014). There are two main approaches in such studies: *marker gene metagenomics*, in which a single gene such as 16S ribosomal RNA is amplified, and *shotgun metagenomics* in which DNA from a sample containing microbes is cleaved into random fragments. With current technologies, amplicon sequencing has much greater sensitivity to low-abundance species, and is therefore popularly used to survey diversity. The diversity in a single sample (*alpha diversity*) is commonly measured using metrics such as the Shannon index (Magurran 1988) and the Chao estimator (Chao 1984), while the variation between pairs of samples (*beta diversity*) is measured using metrics such as the Jaccard distance (Jaccard 1912) or Bray-Curtis dissimilarity (Bray & Curtis 1957). Many such metrics, including Shannon, Chao, Jaccard and Bray-Curtis, are calculated from estimated species frequencies; for further discussion of frequency-based metrics see (Chao et al. 2010). If species cannot be reliably identified or have not been classified by taxonomists, observed traits can be used to define *Operational Taxonomic Units* (OTUs) as proxies for species or higher ranks (Sneath & Sokal 1962). In 16S sequencing, the *de facto* standard approach is to use OTUs constructed as clusters of reads at 97% identity; frequencies are determined as the fraction of reads assigned to each OTU. It is often assumed that OTUs are good approximations to species and that read frequencies are good approximations to species abundances. Despite their central role in typical analyses, these assumptions are rarely stated, questioned or tested in the literature.

Several known factors cause read frequencies to diverge substantially from species frequencies. Prokaryotic genomes contain varying numbers of 16S genes ranging from one to ten or more (Fig. 1), and strains with more genes therefore tend to be more common in the reads (Kembel et al. 2012; Větrovský & Baldrian 2013). PCR amplification efficiency is strongly degraded if a template has mismatches with the primers, causing the number of reads to be suppressed, typically by an order of magnitude or more for each mismatched position (Sipos et al. 2007). With the currently popular V4 primers, ~9% of species have one or more mismatches (Fig. 2). GC content and homopolymers affect polymerase efficiency (Benita et al. 2003; Pinto & Raskin 2012; Strien et al. 2013; Dabney & Meyer 2012). Shorter sequences amplify more efficiently (Dabney & Meyer 2012). When degenerate primers are used, as is commonly the case in 16S sequencing (see e.g. Table 2), biases occur due to unevenness in the oligonucleotide mixture (Polz & Cavanaugh 1998; Kanagawa 2003). Small biases in efficiency are exponentially amplified by the PCR reaction, leading to large biases in read counts (Pinto & Raskin 2012).

**Table 1.** Strains in the HMP mock community. *Assembly* is the Genbank accession for the genome, *Copy nr*. is the number of 16S genes found by SEARCH_16S in the genome, *Primer mism*. notes cases where there are primer mismatches.

| Strain | Assembly | Copy nr. | Primer mism. |
| --- | --- | --- | --- |
| Acinetobacter baumannii, strain 5377 | GCA_000015425.1 | 5 |  |
| Actinomyces odontolyticus, strain 1A.21 | GCA_000154225.1 | 2 |  |
| Bacillus cereus, strain NRS 248 | GCA_000008005.1 | 12 |  |
| Bacteroides vulgatus, strain ATCC-8482 | GCA_000012825.1 | 7 |  |
| Clostridium beijerinckii, strain NCIMB 8052 | GCA_000016965.1 | 14 |  |
| Deinococcus radiodurans, strain R1 (smooth) | GCA_001638825.1 | 3 |  |
| Enterococcus faecalis, strain OG1RF | GCA_000172575.2 | 4 |  |
| Escherichia coli, strain K12, substrain MG1655 | GCA_001566335.1 | 7 |  |
| Helicobacter pylori, strain 26695 | GCA_000307795.1 | 2 | V4-V5 = 1 |
| Lactobacillus gasseri, strain 63 AM | GCA_000014425.1 | 6 |  |
| Listeria monocytogenes, strain EGDe | GCA_000196035.1 | 6 |  |
| Neisseria meningitidis, strain MC58 | GCA_000008805.1 | 4 |  |
| Porphyromonas gingivalis, strain 2561 | GCA_000010505.1 | 4 |  |
| Propionibacterium acnes, strain KPA171202 | GCA_000008345.1 | 3 | V3-V4 = 1, V4 = 2 |
| Pseudomonas aeruginosa, strain PAO1-LAC | GCA_000006765.1 | 4 |  |
| Rhodobacter sphaeroides, strain ATH 2.4.1 | GCA_000273405.1 | 3 |  |
| Staphylococcus aureus, strain TCH1516 | GCA_000017085.1 | 5 |  |
| Staphylococcus epidermidis, FDA strain PCI 1200 | GCA_000007645.1 | 5 |  |
| Streptococcus agalactiae, strain 2603 V/R | GCA_000007265.1 | 7 |  |
| Streptococcus mutans, strain UA159 | GCA_000007465.2 | 5 |  |
| Streptococcus pneumoniae, strain TIGR4 | GCA_000006885.1 | 4 |  |

**Table 2.**
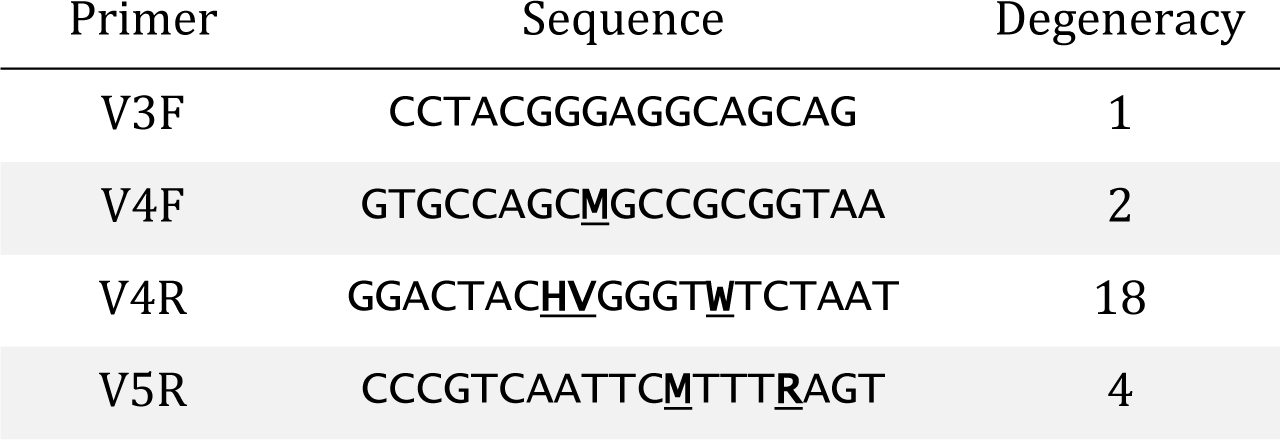
16S primers used by Kozich et al. Wildcard (degenerate) positions are underlined: M={AC}, H={ACT}, W={AT}, R={AG}, V={ACG}. These primers create amplicons in three combinations: V3F-V4R, V4F-V4R and V4F-V5R. All primers except V3F are degenerate, i.e. represent a mixture of different oligonucleotides. Uneven mixing of primers is one of several causes of amplification bias.

**Fig. 1.**
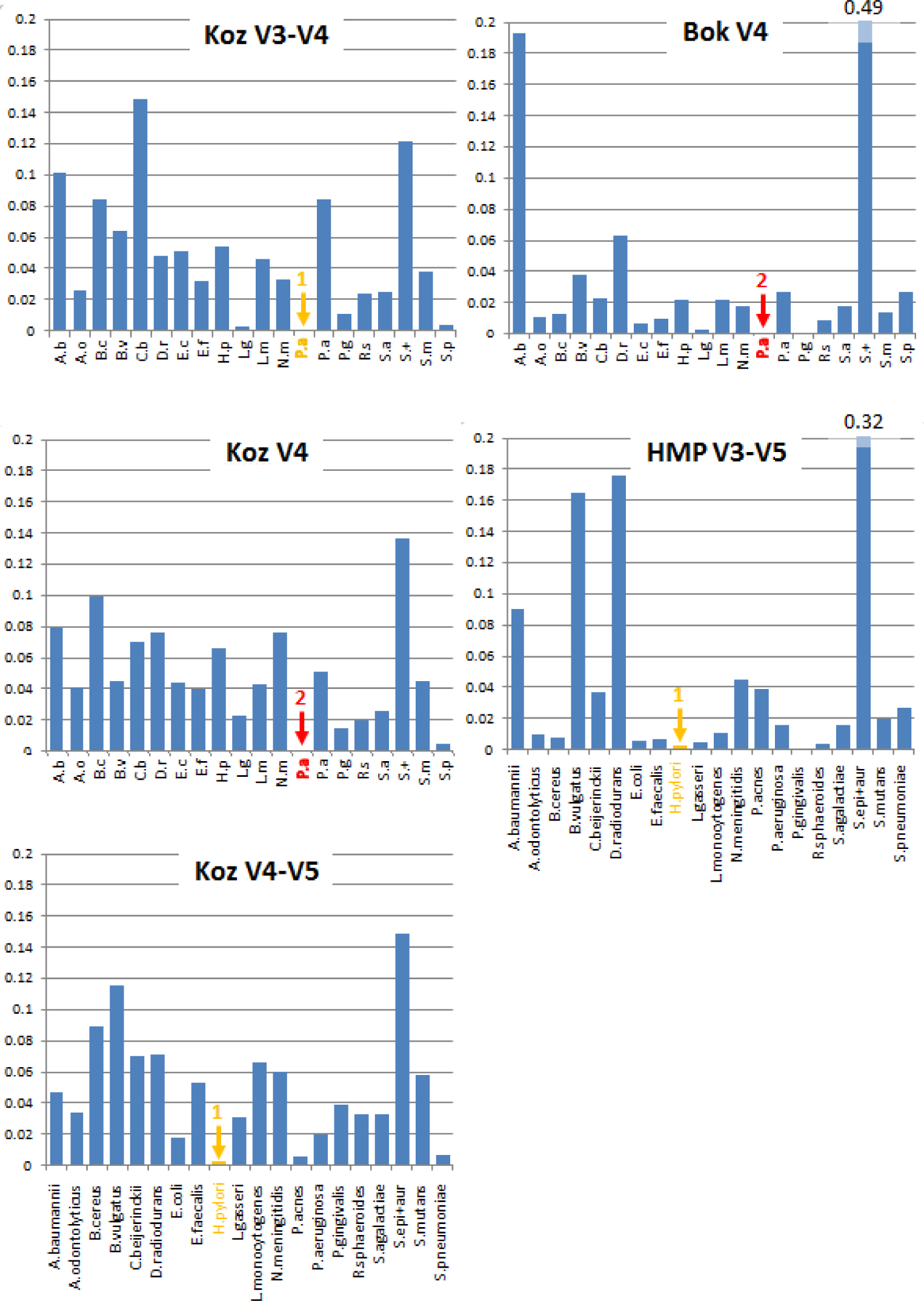
Read abundances for “Even” mock samples. Histograms give measured read frequencies for strains in the mock community (Table 1). Counts for *S. epidermidis* and *S. aureus* are combined into a single pseudo-species *S. epi+aur* because they cannot be reliably distinguished (identical in V4 and V4-V5, one difference in V3-V4). Studies are: *Koz* (Kozich et al. 2013), *Bok* (Bokulich et al. 2013) and *HMP* (HMP Consortium, 2012). The composition of the Even sample was designed to have equal abundances of 16S genes for each species based on the known copy numbers, but the read abundances are in fact highly uneven. Species with primer mismatches are indicated in orange (one difference) and red (two differences), respectively.

**Fig. 2.**
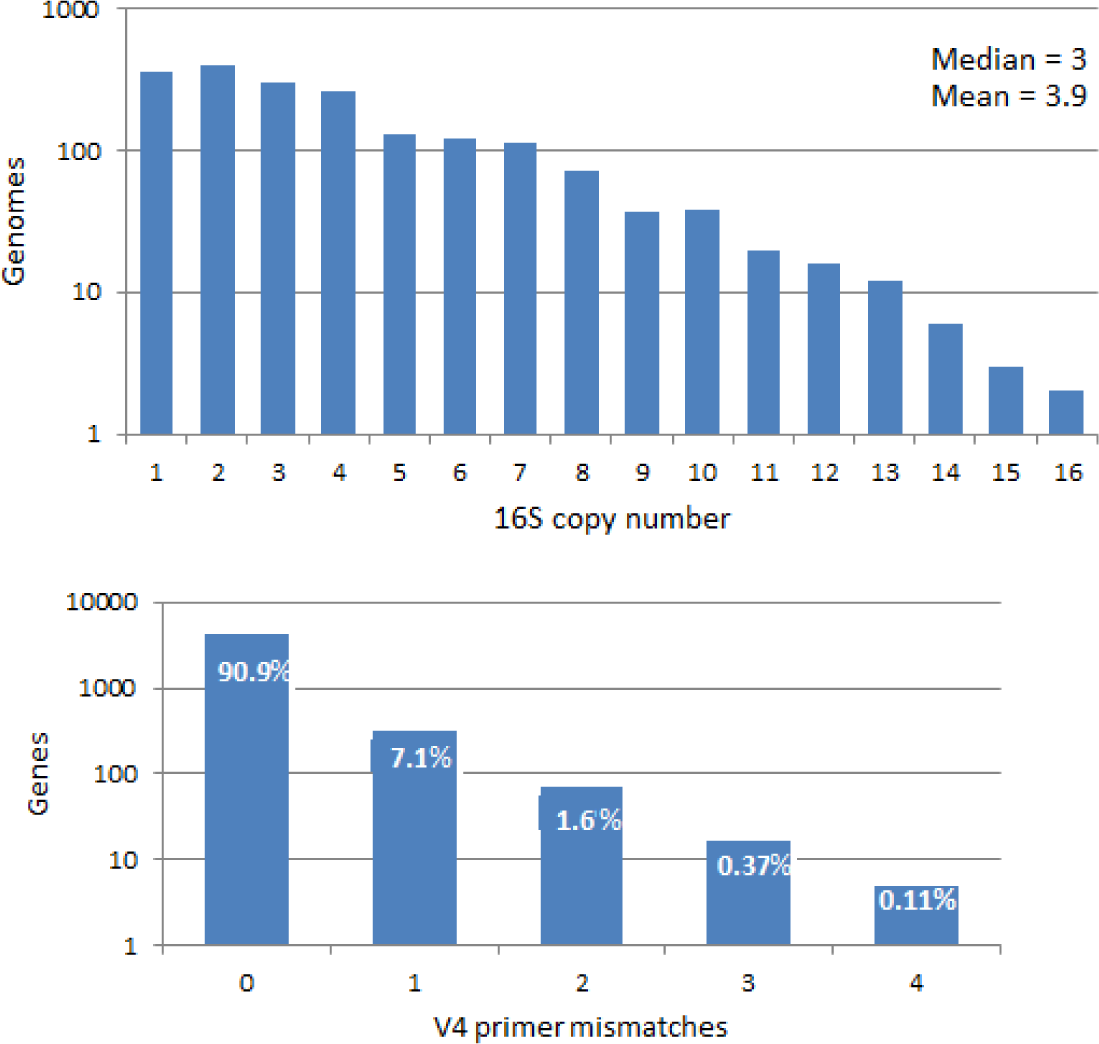
Distributions of 16S copy numbers and V4 primer mismatches in BEST97. The top histogram shows the distribution over 16S copy numbers and the bottom histogram shows the distribution of number of matches with the V4F-V4R primer pair.

An “Even” mock community was developed by the Human Microbiome Project (HMP Consortium 2012) to validate its pyrosequencing protocol. This community has been used as a control sample in several other studies including those of Kozich et al. 2013 (Koz) and Bokulich et al. 2013 (Bok), both of which employed Illumina paired-end sequencing. It is a mix of 21 prokaryotic strains (Table 1) with concentrations designed to give equal mass of 16S genomic DNA from each strain, taking into account the known 16S copy numbers. Naively, each strain should therefore have an approximately equal 16S amplicon read abundance. However, the measured abundances are found to be highly uneven (Fig. 1). Here, I investigate the causes of this disparity. I also describe and validate UNBIAS, an algorithm designed to correct for biases caused by 16S copy number and primer mismatches. At least one previous method has been proposed to correct for copy number bias (Kembel et al. 2012), but to the best of my knowledge no previously published algorithm has attempted to correct for primer mismatch bias. UNBIAS achieves a substantial improvement when the strains are well known, confirming that these biases are significant in practice. However, UNBIAS is less successful when reference sequences have 97% identity or less, as will often be the case, limiting the value of UNBIAS as a practical solution to the problem of abundance bias.

## Methods

### Measuring species abundances from shotgun reads

The Koz data includes shotgun reads of the same mock sample used in 16S sequencing, enabling an accurate measurement of the true genome abundances for each strain. Whole-genome sequences were obtained from the Genbank assemblies given in Table 1. Reads were mapped to these genomes using UBLAST (Edgar 2010), requiring an alignment with ≥98% identity covering ≥90% of the read. The read depth of a genome was calculated as the total number of bases mapped to that genome divided by the genome length. Naively, read depth should be proportional to the genome abundance in the sample. However, regions with high or low GC content are known to be under-represented in shotgun reads (Chouvarine et al. 2016). I therefore calculated GC-corrected depths by selecting reads with GC content in the range 45% to 55% and calculated the mean depth for the subset of bases mapped in those reads (Fig. 3).

**Fig. 3.**
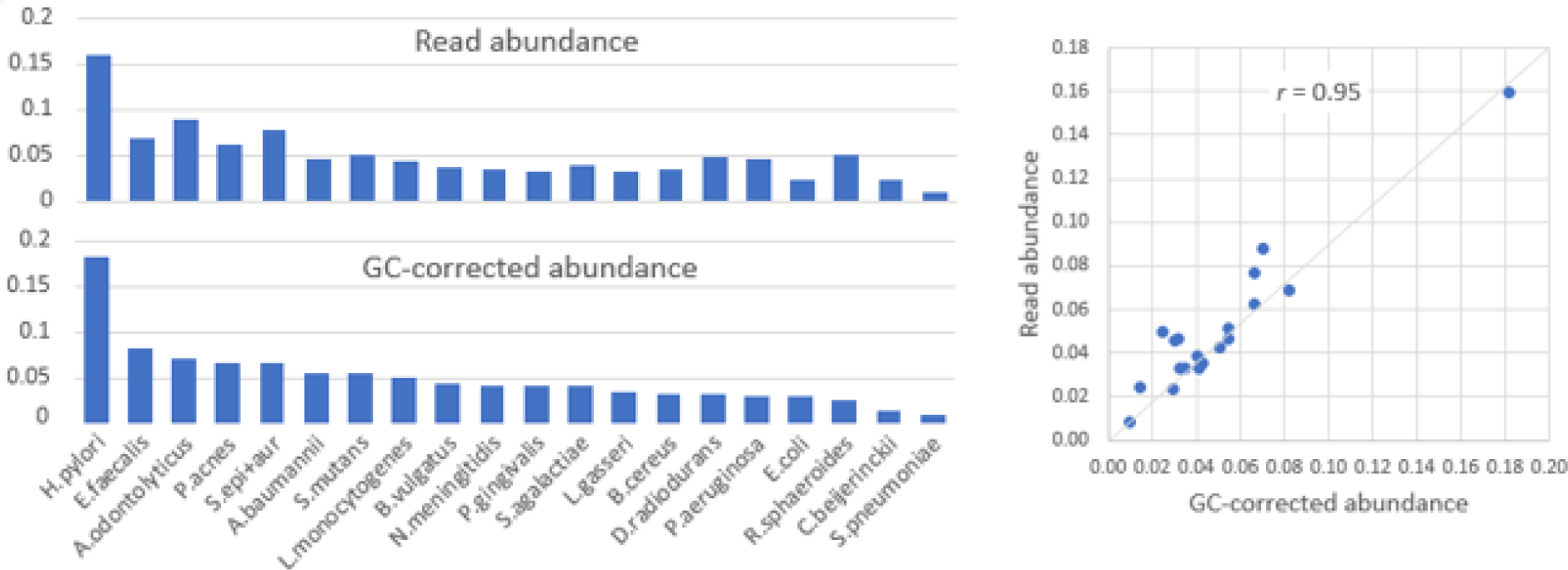
Species frequencies determined by GC-corrected shotgun read abundances. The top histogram shows the abundance of each species calculated from all shotgun reads, below are abundances calculated from reads with GC content between 45% and 55%, which here are taken to be the best estimate of the true frequencies. The scatterplot shows there is good correlation between the raw and corrected frequencies (*r* = 0.95), indicating that shotgun GC bias is a minor issue compared to 16S amplicon sequencing bias.

### BEST and BEST97 databases

The BEST database was constructed as follows. 16S sequences in the 6,487 prokaryotic Genbank assemblies annotated as “complete” (Jan. 2017) were identified using SEARCH_16S (Edgar 2017a), giving a total of 10,849 unique 16S sequences. The BEST97 database contains 1,718 sequences obtained by clustering BEST at 97% identity using UCLUST (Edgar 2010) to compensate for species that have been sequenced many times, e.g. *S. pneumoniae* has 33 complete genomes and *E. coli* has 275.

### UNBIAS algorithm

The UNBIAS algorithm attempts to correct for the two sources of bias I believe to be most important in practice: 16S copy number and primer mismatches. This requires predicting the copy number and mismatch number for each OTU sequence, then adjusting the read count accordingly.

#### Predicting copy number and primer mismatches

Prediction of copy number and primer mismatches is done by the SINAPS algorithm (Edgar 2017b). SINAPS finds the top hit in a reference database using *k*-mer similarity with a confidence estimated obtained by bootstrapping. In each bootstrap iteration, a subset of *k*-mers is selected and used to find the top hit and the trait of interest (here, copy number or primer mismatches) is taken from reference sequence annotation. The trait with highest bootstrap frequency is reported as the prediction, and the frequency with which it occurred is reported the bootstrap confidence. UNBIAS needs a prediction for every OTU, so the bootstrap confidence is ignored. In this case, SINAPS is effectively equivalent to finding the top database hit using the USEARCH algorithm (Edgar 2010).

#### Copy number correction

If the predicted 16S copy number is *C*, the read count is divided by *C*.

#### Primer mismatch correction

If the predicted number of primer mismatches is *m*, the read count is multiplied by 10^*m*^; i.e., an order of magnitude loss in efficiency is assumed for each mismatch. Using 10 as a base is a rather arbitrary choice that probably does not work very well in practice because the true efficiency loss depends on several factors which are unknown or hard to predict. For example, the loss will depend on the mismatch position in the oligonucleotide (mismatches close to the 3' end give higher losses) and will tend to be greater if more rounds of PCR are used. With the Koz data, I found that using 30^*m*^ rather than 10^*m*^ gave better results, typically increasing the correlation by around 0.05 to 0.1, but I felt that tuning this parameter would be misleading as there are only three training examples (*P. acnes* V3-V4, *P. acnes* V4 and *H. pylori* V4-V5; Table 1), and regardless parameters tuned to Koz would be unlikely to generalize well to other datasets using different PCR protocols.

### Idealized UNBIAS

I implemented an idealized variant of UNBIAS in which known copy numbers and primer mismatches are used in place of predictions. This method can of course only be applied to samples of known composition. Idealized results enable assessment of the reduction in correction accuracy due to imperfect predictions and give an upper bound on the accuracy that could be achieved with improved predictions.

## Reduced reference databases

Currently (Jan. 2017) Genbank contains complete genomes for 2,627 different prokaryotic species. These account for only a tiny fraction of extant species (Yarza et al. 2014), and reference databases which annotate 16S sequences with directly measured (as opposed to predicted) traits such as 16S copy number and taxonomy are therefore very sparse. This can be quantified by considering the identity distribution between the OTU sequences in a sample and a reference database (Fig. 8). Strains in a mock community are usually well-known and are therefore likely to be present in any given reference database. For example, all strains in the HMP mock community have finished genomes (Table 1). Using a mock sample with a database based on all complete genomes is therefore an unrealistic test of UNBIAS. To address this, I created reduced reference databases by discarding all 16S sequences above a given identity (97%, 95% or 90%) with the mock sequences. Identities below 90% are often encountered in practice (Fig. 8), but this is already below the limit where UNBIAS can provide a useful correction (Fig. 6).

**Fig 4.**
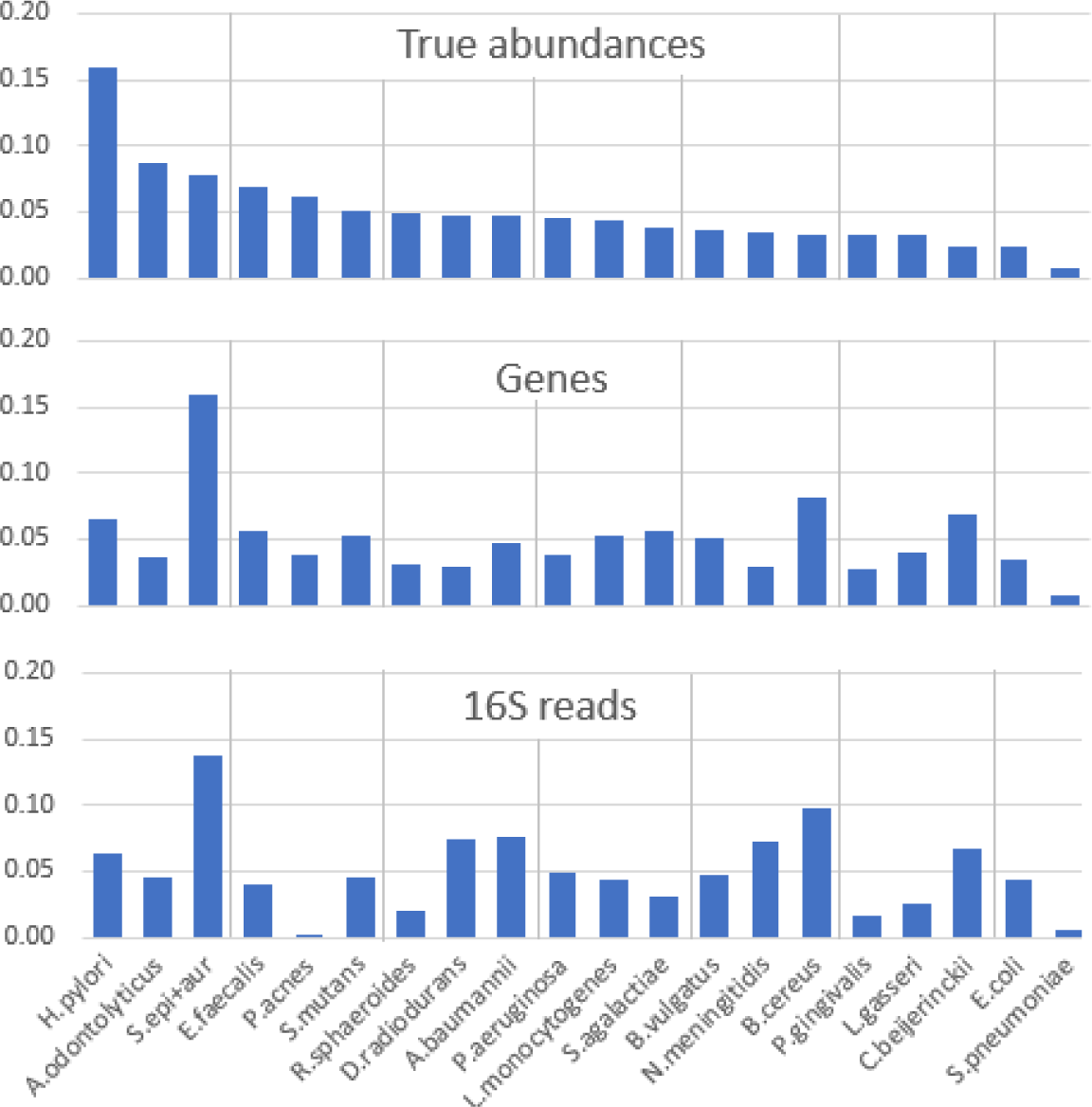
Abundances for all Even samples in the Koz V4 data. Top are the true abundances calculated from GC-corrected shotgun read depths (Fig. 3). Middle are the gene abundances obtained by multiplying the true abundances by the known 16S copy numbers. By design of the Even community, the gene abundances should be uniform; divergences from uniform frequencies are interpreted as imperfect mixing of the strains when the sample was prepared. Bottom are abundances obtained from the 16S amplicon reads. Divergence between gene abundances and 16S read abundances are interpreted as PCR and sequencing bias.

**Fig. 5.**
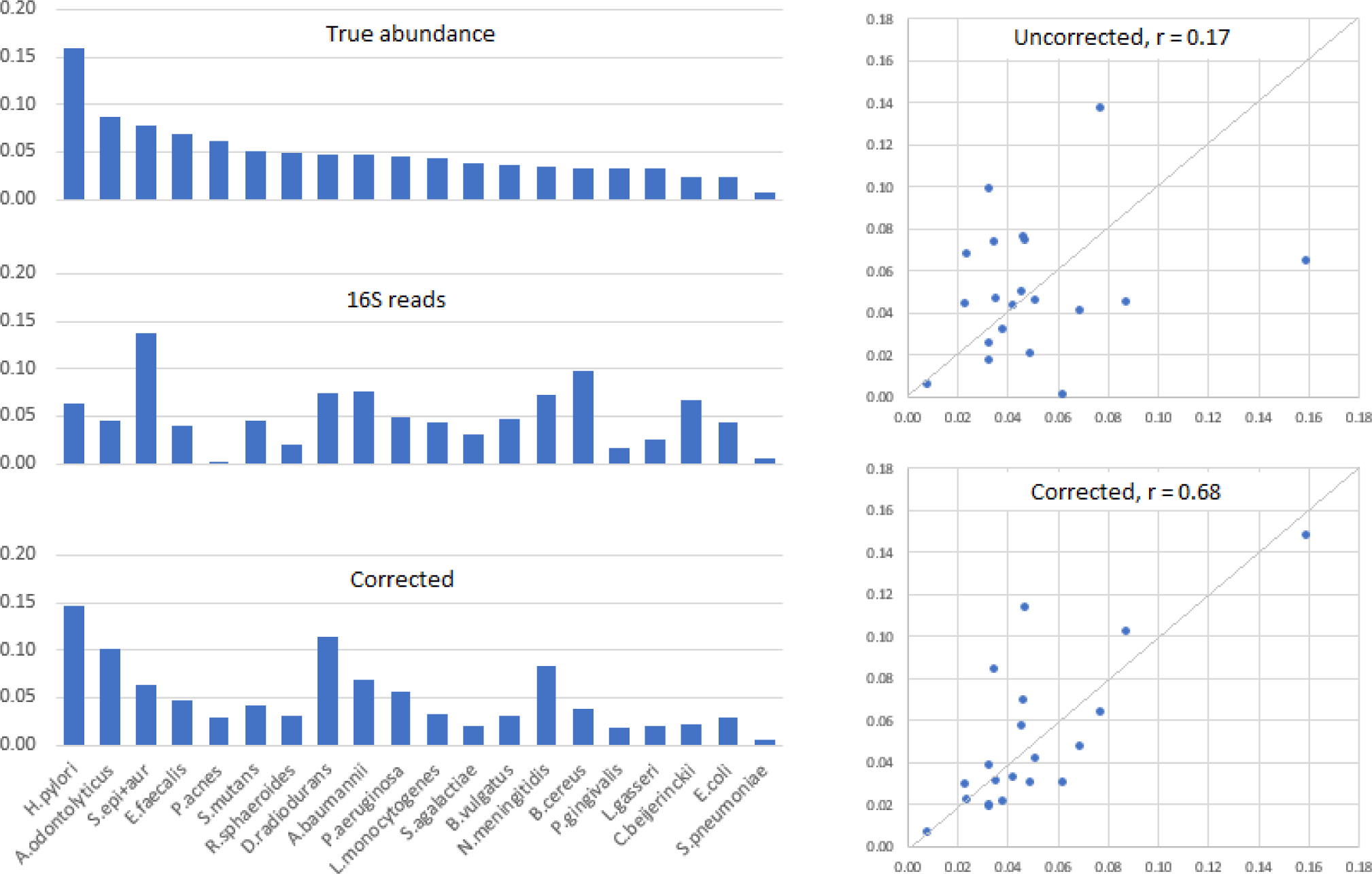
V4 read abundances before and after correction by idealized UNBIAS. Top left: true genome abundances measured from shotgun reads. Middle left: 16S read abundances. Bottom left: frequencies estimated by UNBIAS. Top-right: scatter plot of read abundances (*y* axis) vs. true abundances (*x*), bottom-right scatter plot of corrected abundances (*y*) vs. true abundances (*x*). The correlation is improved from *r* = 0.17 to *r* = 0.68; however, this is unrealistic because the UNBIAS estimates were idealized, i.e. based on known rather than predicted copy numbers and primer mismatches. Also, read abundances were averaged over all 33 replicates in the 11 Koz runs. Single samples exhibit fluctuations and the correlation when idealized UNBIAS is applied to single samples is therefore usually lower (mean 0.60, see Fig. 6).

**Fig. 6.**
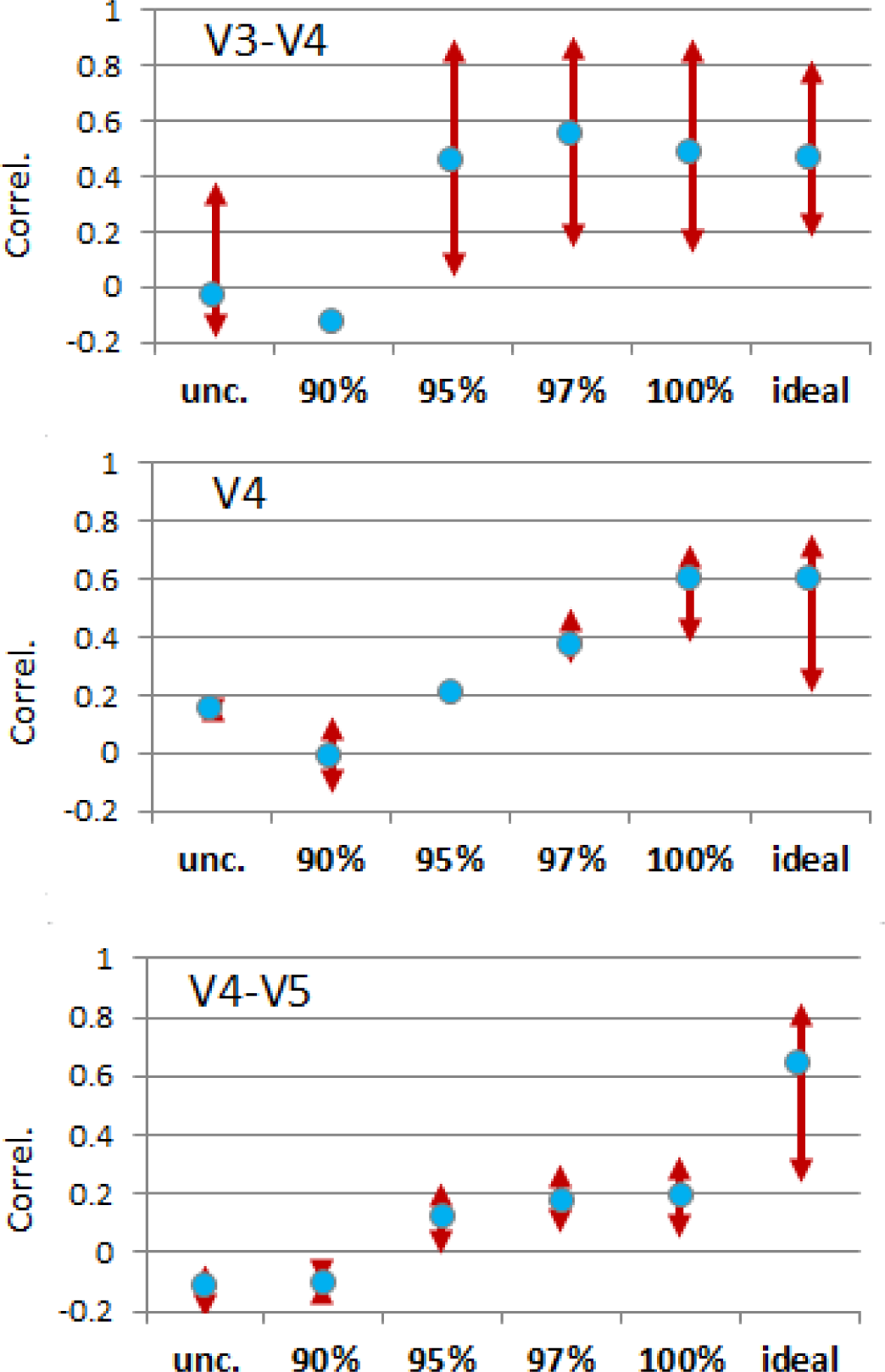
Correlation between estimated and true species abundances. The *y* axis is the Pearson correlation coefficient *r*. Results are shown for the uncorrected read frequencies (*unc*.), for a best-case scenario where the copy numbers and primer mismatches are known (*ideal*) and for identities 90%, 95%, 97% and 100% between the 16S sequences and the closest reference sequence. The 100% identity case gives different results from the ideal case because the sequence of a given tag may be identical to two or more reference species, as with the V4 segments of *S. epidermidis* and *S. aureus*. Blue dots show the mean value; the red arrows show the range from minimum to maximum *r* over the 33 replicate samples.

**Fig. 7.**
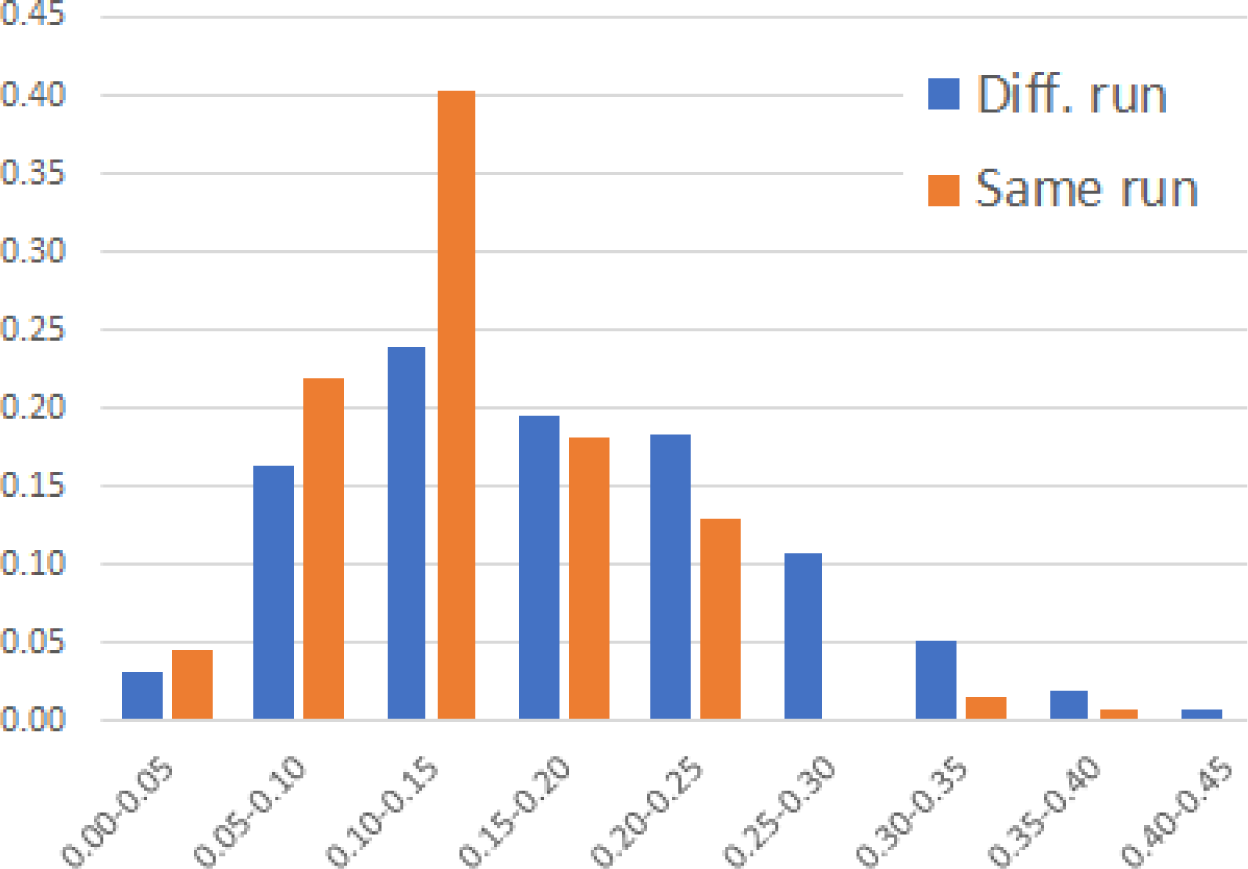
Jaccard distances for pairs of replicate samples. For each pair of identical samples sequenced in replicate, the Jaccard distance was calculated and binned into intervals of 0.05. Blue bars (first for each bin) show the distribution over pairs sequenced in different runs, orange bars (second) the distribution over pairs sequenced in the same run. The *x* axis is the frequency normalized to sum to 1.0 over all bins.

**Fig. 8.**
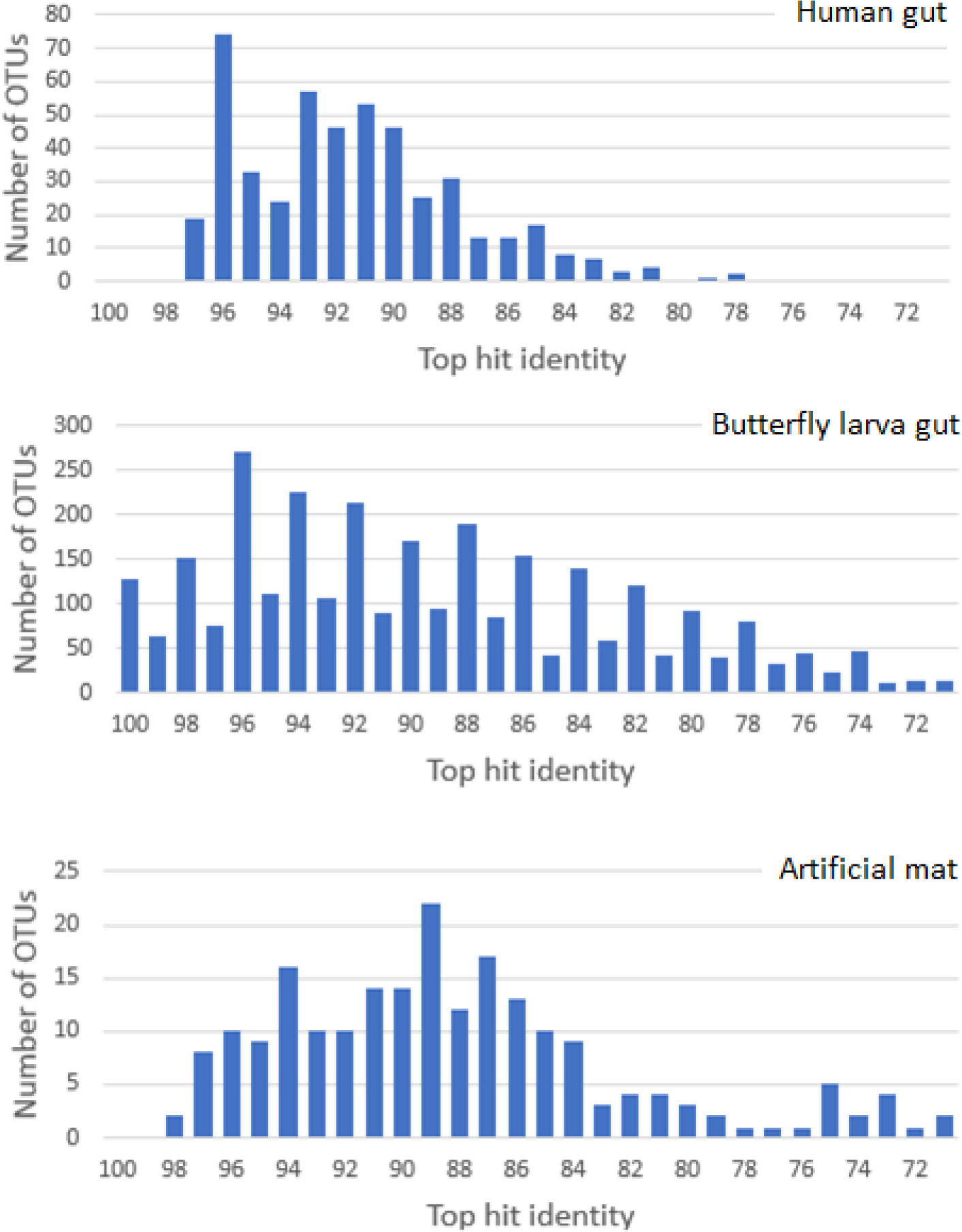
16S identities of OTUs with finished genomes. The figure shows the distribution of top hit identities between UPARSE OTUs from representative projects PRJEB16057, PRJEB17686 and PRJEB14970 in the NCBI Short Read Archive with BEST (16S sequences from complete genomes) as a reference. PRJEB16057 contains human gut samples, PRJEB17686 is butterfly larva gut and PRJEB14970 is artificial microbial mats.

## Measuring correlation

Correlations were measured using the Pearson correlation coefficient (*r*) (Pearson 2006). For two frequency distributions ***x***, ***y*** with *x_i_, y_i_, i* = 1… *N, r* is calculated as

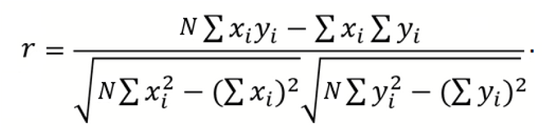
The correlation is measured by finding a line that minimizes the root mean square distance of (*x, y*) points from the line. If *r* is 1 or −1, there is an exact linear relationship between ***x*** and ***y***; i.e., all (*x, y*) points fall on the line. If *r* < 0, the best line has a negative slope, otherwise the slope is positive. If *r* is close to zero, the best line is a poor fit to the points, indicating that the relationship is not linear.

## Comparing species frequencies (beta diversity)

Frequency distributions ***x**, **y*** of two samples were compared using the Jaccard distance (*J*) (Jaccard 1912), calculated as

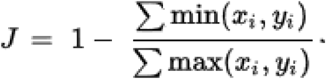
If *J* is 1, then no OTU is present in both ***x*** and ***y*** (equivalently, every OTU is found in only one of the samples), indicating that the samples are maximally different. If ***x*** and ***y*** are identical, *J* is zero. If the same sample is sequenced in two replicates, we would therefore ideally find that *J* = 0, but in practice *J* will be greater than zero due to cross-talk (Kircher et al. 2012; Nelson et al. 2014; Edgar 2017c), fluctuations caused by selection effects, etc.

## Modeling the impact of bias

To investigate the impact of bias in practice, I implemented BIASIM, a simple simulation of bias implemented as a short Python script (available on request). I used the geometric series model (Motomura 1932) where the species with rank *k* > 0 has abundance *f_k_* given by

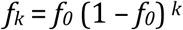
where *f_0_* is the frequency of the first (dominant) species. The value of *f_0_* is selected from the range *F_min_* to *F_max_* with a uniform probability distribution, and frequencies for *N* species are generated. For this work, I used the following parameters *F_min_* = 0.1, *F_max_* = 0.5, *N* = 100. Then, the distribution is biased by choosing a copy number and primer mismatch number at random for each species. The copy number *C* is chosen from the range 1 to 8 with uniform probability. The primer mismatch number *d* is 0 with probability 0.9, 1 with probability 0.09 and 2 with probability 0.01. The biased abundance *f_k_′* is calculated as

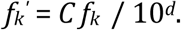

## Results

### Distribution of 16S copy number

Fig. 2 (top histogram) shows the distribution of 16S copy number for 16S sequences in the BEST97 database. The median copy number is 3, with a mean of 3.9. Copy numbers from 1 to 10 are common, i.e. a range of an order of magnitude, indicating that copy number bias is likely to be substantial in practice.

### Distribution of V4 primer mismatch number

Fig. 2 (bottom histogram) shows the distribution of V4 primer mismatches for sequences in BEST97. This shows that ~9%, i.e. about one in ten, species has at least one mismatch with the currently popular V4 primer pair used for Illumina 2×250 sequencing. The HMP mock community has 21 strains. Using 9% as an estimate, we expect that 21×9% = 1.9 of these strains will have mismatches, and in fact we find that one (*P. acnes*) has two V4 mismatches (Table 1). If BEST97 is representative of species encountered in practice, then we can expect that most species, about 9 in 10, are unaffected by primer mismatch bias. The remaining one in ten will be strongly affected because the efficiency loss is at least a factor of 10 per mismatch.

### Can we infer the dominant species from 16S reads?

How important are these biases in practice? Of course, this depends on which biological question you want to answer. As an example, can you infer the dominant (most abundant) species from the reads? To investigate this, I used BIASIM (described in Methods) to generate 1,000 simulated communities. I found that the mean rank of the dominant species in the reads was 3.0 and the most abundant species in the reads was different from the true dominant species in 62% of the communities. While of course BIASIM is a simplified model that would be inadequate in some contexts—for example, the geometric series model is not a good fit to many communities (Neuteboom & Struik 2005)—the details should not matter much for this purpose and I believe it is sufficiently realistic to give a credible estimate. The Koz data provides an illustrative example. *H. pylori* is the dominant species with an abundance ~2x the second-ranked species (*A. odontolyticus*), but its rank is 6th in the V3-V4 reads, 7th in the V4 reads, and is 20th (last) in the V4-V5 reads, where it has a primer mismatch. The low ranks found in V3-V4 and V4 can be explained because *H. pylori* has only two 16S genes, fewer than all the other strains except *A. odontolyticus* which also has two (Table 1).

### Performance of idealized UNBIAS

Fig. 5 shows the improvement in correlation achieved by idealized UNBIAS over all Even samples in Koz. The correlation is improved from *r* = 0.17 to *r* = 0.68. Comparing the histograms at top-left and bottom-left, we can see that the corrected distribution is clearly better, but still substantially different from the true distribution. However, this result is unrealistic because copy number and primer mismatches are known rather than predicted, and because the read abundances are averaged over many samples (three replicates in each of 11 runs, for a total of 33 samples).

### Performance of UNBIAS on individual samples

Fig. 6 shows the range of correlations for uncorrected (*unc*.) and corrected abundances, processing each of the 33 samples individually. Reads were sub-sampled to a total of 5,000 reads per sample to mitigate the effects of cross-talk, which causes many spurious low-abundance OTUs in this data (Edgar 2017c). *Ideal* is the idealized case where known rather than predicted copy numbers and mismatch numbers are used. Other ranges marked 90%, 95%, 97% and 100% give results using the full (100%) and reduced (90% to 97%) reference databases to assess performance when OTU sequences are diverged from the reference database (see Fig. 8). These results show that correlations of ~0.5 to 0.6 are achieved when sequences are present in the database but fall to close to zero at 90% identity. Fig. 8 shows that OTUs with <90% identity are common, which indicates that UNBIAS has limited value as a practical tool.

### Variations in replicate samples

Fig. 7 shows the range of Jaccard distances for all pairs of sub-sampled mock community samples in the Koz data (11 runs, 3 replicate samples per run). This shows that the mean Jaccard distance is ~0.15 for samples in the same or different runs with high frequencies for values up to 0.25 and a tail extending to around 0.45. The maximum value was 0.71. The mean frequency correlation between replicates was 0.82 with a standard deviation of 0.17. Variations between replicates must be due to unpredictable rather than systematic effects. If we assume that replicate variations seen in the Koz data are typical of other experiments, this shows that ~0.8 is an approximate upper bound on the correlation between measured and true abundances that could be achieved by an ideal algorithm that achieved the best possible correction for systematic biases (e.g., if all species were fully characterized by a reference database).

## Conclusions

The results presented here show that systematic bias due to 16S copy number and primer mismatches cause read abundance distributions to be substantially different from the true species abundance distributions. Primer mismatches are relatively easy to predict because a full-length sequence suffices, and databases such as SILVA, RDP and Greengenes have ~10^6^ full-length 16S sequences. However, measurement of copy number usually requires a whole-genome sequence, which is not available for the vast majority of extant microbial species. While it is possible to predict copy number when OTUs have sufficiently high identity with known genomes (Kembel et al. 2012; Edgar 2017b), this is not sufficient to infer a useful approximation to the species abundance distribution in a typical experiment where OTU identities are often too low with known reference sequences. An identity of 90% corresponds roughly to family level (Yarza et al. 2014), and copy number is not well enough conserved at this distance to enable accurate predictions (Edgar 2017b). The underlying problem is therefore sparse reference data with insufficient coverage, which cannot be addressed by using another type of prediction algorithm (say, ancestral reconstruction). The difficulties of predicting copy number and primer mismatches are compounded by unpredictable intra-sample variations which cause correlations of ~0.8 and Jaccard distances of ~0.15 between replicates of identical samples in a representative dataset.

In conclusion, 16S read abundances have no useful correlation with species abundances in a single sample, and this problem seems likely to remain unsolved unless new sequencing methods are developed or a dramatic increase in coverage of whole-genome sequences is achieved. It may nevertheless be possible to measure between-sample variations (beta diversity) when they exceed intra-sample variations between replicates, but the requirement to establish appropriate thresholds to distinguish between biological variation and experimental variation is rarely, if ever, considered. These observations call into question the naive use of alpha and beta diversity metrics which implicitly assume that read abundances are a good approximation to species abundances.

